# Fear extinction requires infralimbic cortex projections to the basolateral amygdala

**DOI:** 10.1101/172791

**Authors:** Daniel W. Bloodgood, Jonathan A. Sugam, Andrew Holmes, Thomas L. Kash

## Abstract

Fear extinction involves the formation of a new memory trace that attenuates fear responses to a conditioned aversive memory, and extinction impairments are implicated in trauma- and stress-related disorders. Previous studies in rodents have found that the infralimbic prefrontal cortex (IL) and its glutamatergic projections to the basolateral amygdala (BLA) and basomedial amygdala (BMA) instruct the formation of fear extinction memories. However, it is unclear whether these pathways are exclusively involved in extinction, or whether other major targets of the IL, such as the nucleus accumbens (NAc) also play a role. To address this outstanding issue, the current study employed a combination of electrophysiological and chemogenetic approaches in mice to interrogate the role of IL-BLA and IL-NAc pathways in extinction. Specifically, we used patch-clamp electrophysiology coupled with retrograde tracing to examine changes in neuronal activity of the IL and prelimbic cortex (PL) projections to both the BLA and NAc following fear extinction. We found that extinction produced a significant increase in the intrinsic excitability of IL-BLA projection neurons, while extinction appeared to reverse fear induced changes in IL-NAc projection neurons. To establish a causal counterpart to these observations, we then used a pathway-specific Designer Receptors Exclusively Activated by Designer Drugs (DREADD) strategy to selectively inhibit PFC-BLA projection neurons during extinction acquisition. Using this approach, we found that DREADD-mediated inhibition of PFC-BLA neurons during extinction acquisition impaired subsequent extinction retrieval. Taken together, our findings provide further evidence for a critical contribution of the IL-BLA neural circuit to fear extinction.

## Introduction

Fear extinction, a process by which learned fear responses are reduced through new inhibitory learning, has emerged as a translationally valuable assay for studying anxiety and trauma-related disorders (1-3). Previous evidence from multiple lines of inquiry strongly implicates the medial prefrontal cortex (mPFC) in the mediation of extinction (1, 4-6), but the precise position of this region within the broader brain network underlying extinction remains incompletely understood. Recent studies in mice have shown that neuronal outputs from the ventromedial PFC (vmPFC), containing the infralimbic cortex (IL) region, to the basolateral (BLA) (7-9) and basomedial (BMA) (10), as well as reciprocal connections from BLA to IL (11, 12) and projections from BLA to NAc (13) are important for extinction.

While these data suggest IL-BLA projection neurons represent a critical circuit for extinction, they require substantiation using alternative approaches. Current data also fail to adequately address the question of whether the functional role of the IL-BLA circuit is redundant to other major IL projection pathways. For example, it remains unclear whether IL projections to the nucleus accumbens (NAc) mediate extinction despite reports that functional manipulations in the NAc, including dopamine D2 receptor blockade (14) and virally-driven activation of the transcription factor cAMP response element binding protein (CREB) (15), can produce significant deficits in fear extinction.

The goal of the current study was to further examine the contribution of the IL-BLA pathway to fear extinction and determine whether IL-NAc projections played a complementary or distinct role in extinction. To this end, we first traced projection neurons from IL and PL to the BLA and NAc, in order to evaluate the degree to which the prefrontal efferents to these regions overlapped. We next performed *ex vivo* electrophysiological recordings to test whether fear extinction differentially affected the excitability of IL-BLA, PL-BLA and IL-NAc projection neurons. To then establish causal contributions of the PFC-BLA to extinction, we chemogenetically inhibited this pathway and assessed the behavioral consequences.

## Materials and Methods

### Subjects

Adult male C57BL/6J mice (The Jackson Laboratories, Bar Harbor, ME, USA), at least 8 weeks of age, were used for all experiments. Mice were group-housed in a colony room with 12:12 h light-dark cycle with lights on at 0700 hr. Mice had *ad libitum* access to rodent chow and water. The Institutional Animal Care and Use Committee of the University of North Carolina at Chapel Hill approved all procedures.

### Neuroanatomical tracing

PFC cells projecting to the BLA or NAc were visualized using the retrograde tracer Cholera Toxin B (CTB) (Invitrogen, ThermoFisher Scientific, Waltham, MA, USA). Using stereotaxic surgery, 0.3 μl CTB labeled with either Alexa 488 or Alexa 555 (0.5% weight/volume, in sterile PBS) was bilaterally delivered into the BLA and NAc at a rate of 0.1 μl per minute using a 1 μl Hamilton Syringe (Hamilton Company, Reno, NV). The dye conjugate (Alexa 488 or Alexa 555) that was injected into each region was counterbalanced across mice (for schematic, see **Figure 1A, upper panel**). The injection coordinates for the BLA were 1.3 mm posterior to bregma, 3.25 mm lateral to the midline and 4.95 mm ventral to the skull surface. The coordinates for the NAc were 1.3 mm anterior to bregma, 0.85 mm lateral to the midline and 4.75 mm ventral to the skull surface. The injection needle was left in place for at least 5 minutes to ensure diffusion.

**Figure 1:**
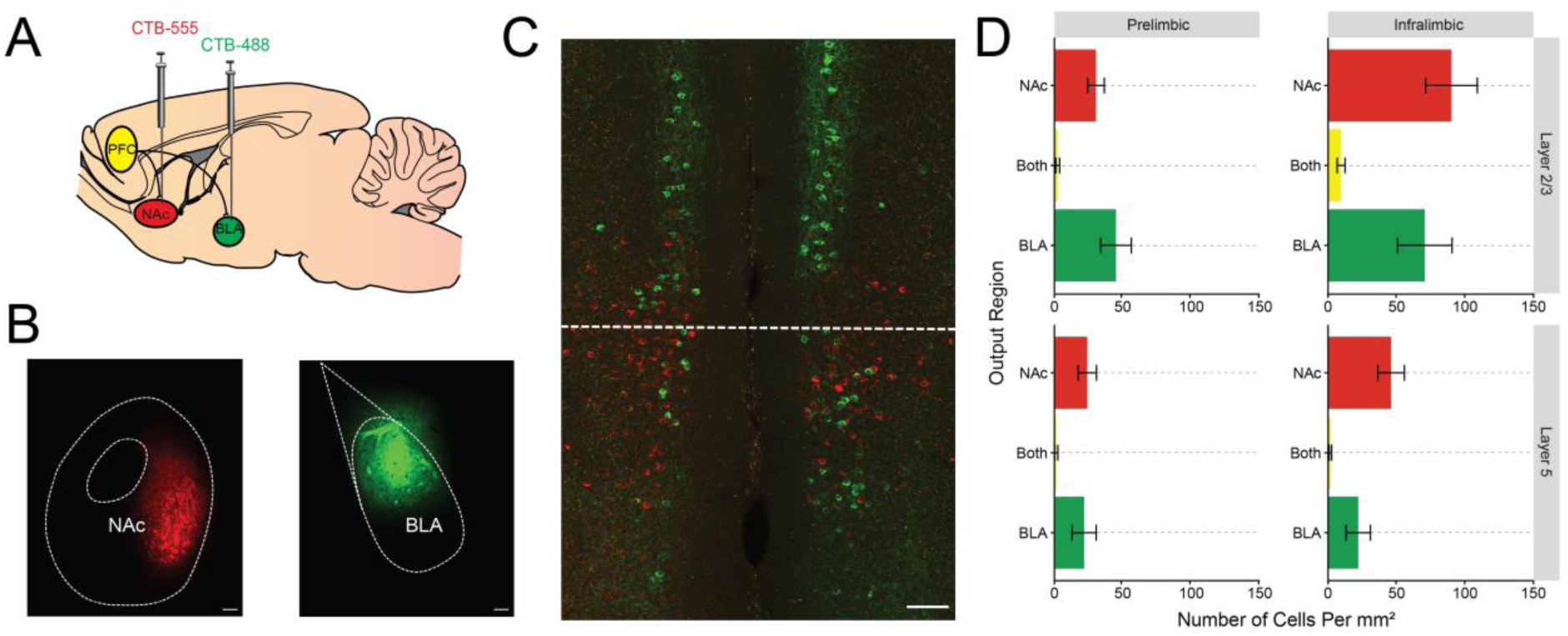
Distinct mPFC projections to the BLA and NAc. (**A**) The retrograde tracer CTB separately labeled with Alexa 488 and Alexa 555 was injected into the NAc and BLA. (**B**) Representative image of injection sites in the BLA and NAc. (**C**) Injection of tracers led to dense, but non-overlapping labeling of cells in the IL and PL. (**D**) Quantification of fluorescent cells revealed dense, nonoverlapping labeling of BLA and NAc outputs throughout the PFC that was concentrated in the superficial layers of IL. Scale bar represents 100 μm. All data presented are mean ± SEM. N=18 images from n=6 mice.

One week after surgery, mice were deeply anesthetized with an overdose of tribromoethanol (250 mg/kg, intraperitoneal) and transcardially perfused with 0.1 M phosphate buffered saline (4°C, pH 7.4) followed by 4% paraformaldehyde in PBS. Brains were post-fixed for 24 hours in 4% PFA and then cryoprotected in 30% sucrose in PBS. Tissue sections containing the PFC, NAc, and BLA were cut into 45 μm sections using a Vibratome (Leica VT1000S, Buffalo Grove, IL, USA) and stored in a 50% glycerol/PBS solution.

Correct placements for CTB infusions were verified using a wide-field epifluorescent microscope (BX-43, Olympus, Waltham, MA, USA) using a stereotaxic atlas (Franklin & Paxinos, 2008). Quantification of CTB fluorescence and representative images of virus expression were collected on a Zeiss 800 Laser Scanning Confocal Microscope (20x objective, NA 0.8) (Carl Zeiss, Jena, Germany). Images were cropped and annotated using Zeiss Zen 2 Blue Edition software (for example, see **Figure 1B**). Regions of Interest (ROIs) corresponding to Layers 2/3 and Layer 5 of IL and PL were drawn based on areas specified in the Allen Mouse Brain Atlas (Allen Institute, Seattle, WA, USA). Quantification of CTB positive cells was performed using manual counts with the cell counter plugin in FIJI (Curtis Rueden, LOCI, University of Wisconsin-Madison, Madison, WI, USA). A preliminary quantification showed that 98.2% of CTB positive cells were also positive for the neuronal marker NeuN, showing that CTB almost exclusively labels neuronal cell types. Thus, DAPI was used as a marker of nuclei and acquired as a separate channel for all the images used for quantification. The DAPI channel has been omitted from the representative image to more clearly show the distribution of CTB positive neurons.

### Behavioral procedures

Mice underwent fear conditioning and extinction as described previously (16). Briefly, fear conditioning was performed in Context A: a 30.5 x 24.1 x 21.0 cm sound-attenuating chamber with metal walls and grid floor, cleaned with 20% ethanol paired with a vanilla scent to provide an olfactory contextual cue. Following a 120-second baseline habituation period, a 30-second 8 kHz 80 dB tone (CS) was presented, co-terminating with a 2-second 0.6 mA footshock (US) delivered through the grid floor. There were 5 x CS-US pairings, interspersed by a 20-120 second pseudorandomized interval and a 120-second post conditioning period after the final pairing. Freezing behavior was measured as an index of fear learning, and was operationally defined as making no movement other than what is necessary to breathe. Freezing behavior was quantified using an automated videotracking software (Ethovision 9.0, Noldus, Leesburg, VA, USA) with a 1-second minimum threshold for the detection of freezing behavior.

The following day, mice underwent extinction acquisition in a novel Context B. The chamber used was identical to Context A, but was modified with a clear circular plastic white surround and a plastic white insert over the grid floor. In between trials, the chamber was cleaned with 70% ethanol without a vanilla scent to provide an olfactory cue that served as another distinct attribute of the novel context. Following a 120-second baseline habituation period, there were 50 x CS presentations interspersed by a 5-second inter-trial interval. For statistical analysis, the responses were binned across five CS presentations to examine changes in freezing behavior across ten cue blocks. In the chemogenetic experiment, there was an additional test day during which mice were returned to Context B the day after extinction acquisition to test for extinction retrieval and, following a 120-second baseline habituation period, presented with 5 x CS presentations interspersed by a 5-second inter-trial interval.

### Slice electrophysiology

IL and PL cells projecting to the BLA or NAc were visually differentiated using fluorescent Retrobeads (for schematic, see **Figure 2A, 3A**). Using the same stereotaxic coordinates and procedures described above, 0.3 μl Red Retrobeads (Lumafluor Inc, Durham, NC, USA) were bilaterally delivered at a rate of 0.1 μl per minute to the BLA or NAc, in different groups of mice. One week after surgery, mice underwent either fear conditioning or extinction training, as described above, and then 10 minutes later, were deeply anesthetized with isoflurane and decapitated. An experimentally-naïve group was also sacrificed. The brain was rapidly removed and placed in ice-cold sucrose-artificial cerebrospinal fluid (aCSF) containing (in mM) 194 sucrose, 20 NaCl, 4.4 KCl, 2 CaCl_2,_ 1 MgCl_2,_ 1.2 NaH_2_PO_4_, 10 glucose and 26 NaHCO_3_. Three hundred micron acute brain slices were sectioned on a vibratome and transferred to a submerged recording chamber (Warner Instruments, Hamden, CT, USA), where they were perfused with heated (30-31 Celsius), oxygenated aCSF at a rate of 2 ml per minute and allowed to equilibrate for 30 minutes before recordings.

**Figure 2:**
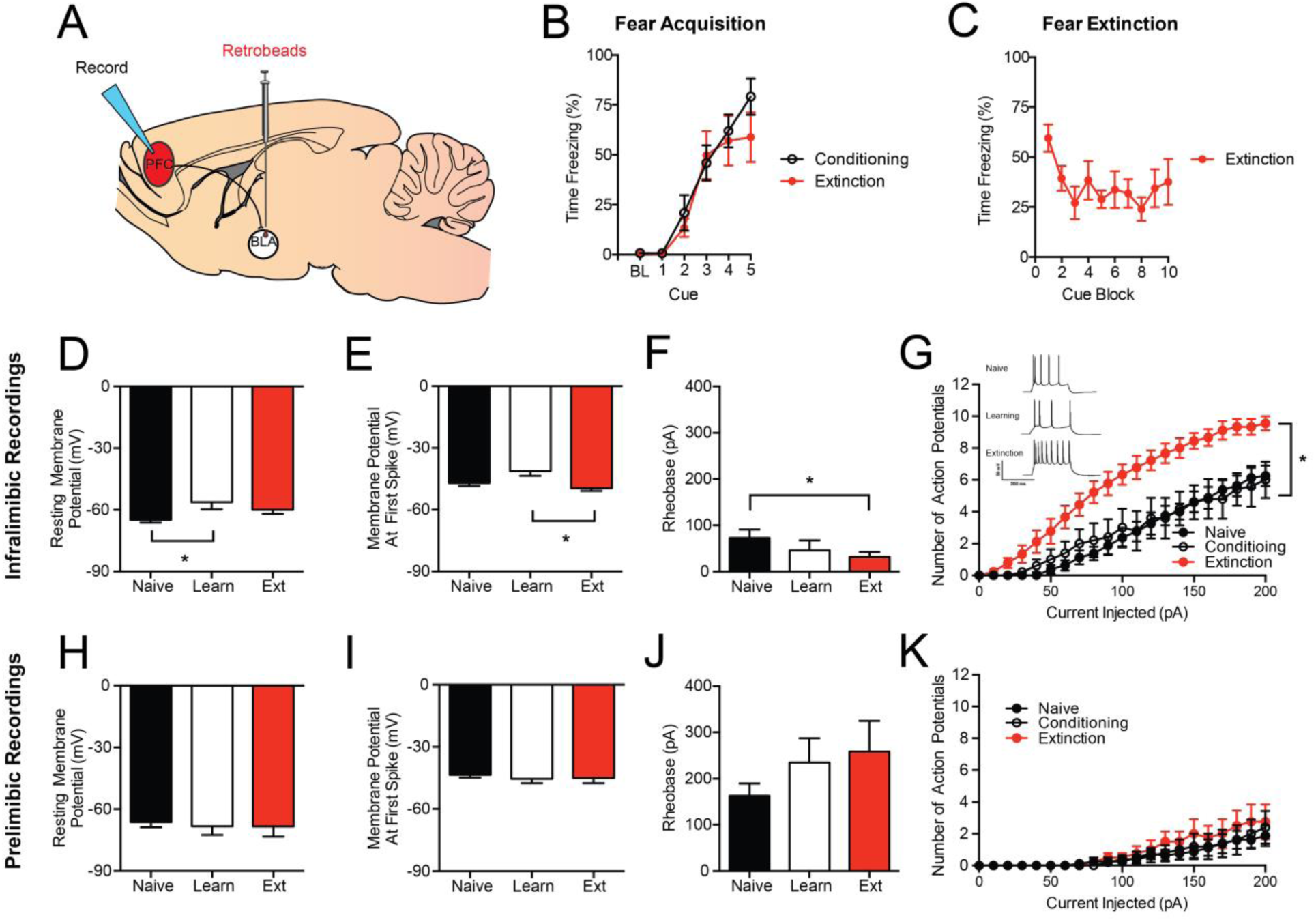
Extinction increases neuronal excitability in the IL-BLA, but not PL-BLA, pathway. (**A**) Retrobeads were injected into the BLA to visualize IL-BLA and PL-BLA projection neurons for electrophysiology recordings. (**B**) Fear conditioning increased freezing across CS-US pairings. (**C**) Fear extinction decreased freezing across CS trial-blocks. In IL-BLA projection neurons, fear conditioning resulted in (**D**) increased resting membrane potential that was not present after extinction. In IL-BLA neurons, following extinction training there was also (**E**) higher action potential threshold, (**F**) lower rheobase, and (**G**) an increase in the number of action potentials fired across increasing current injections. (Inset) Representative traces of current injected firing during the final step of an increasing current injection protocol. In PL-BLA projections neurons, there were no differences in (**H**) resting membrane potential, (**I**) action potential threshold, (**J**) rheobase, or (**K**) action potential number with increasing current injection magnitude. Data presented are tests completed at resting membrane potential. All data presented are mean ± SEM. N=6 mice per group, n=7-9 cells per group, per brain region. * denotes p<0.05.

**Figure 3:**
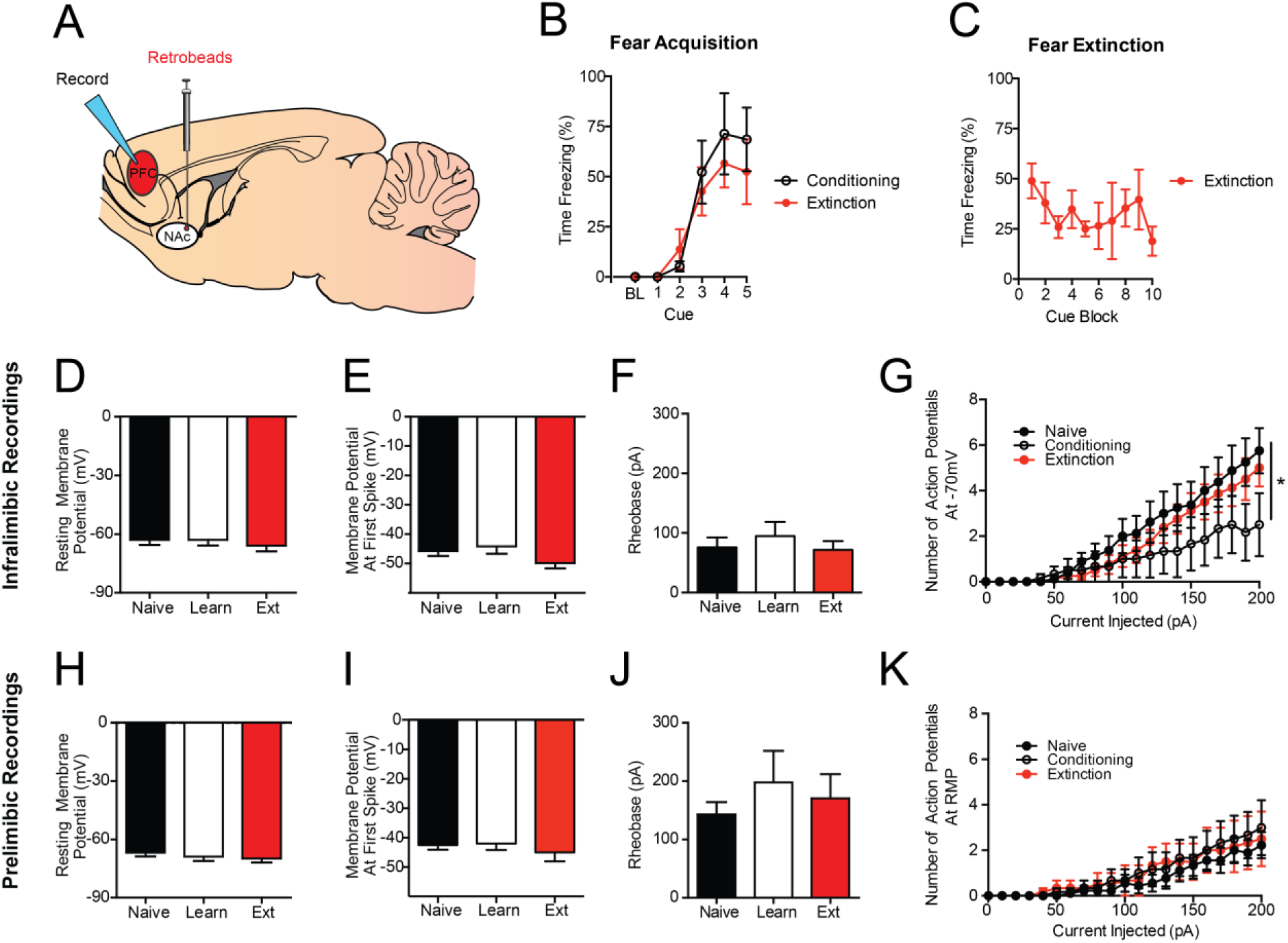
Extinction does not increase neuronal excitability in the IL-NAc or PL-NAc pathways. (**A**) Retrobeads were injected into the NAc to visualize IL-NAc projection neurons for electrophysiology recordings. (**B**) Fear conditioning increased freezing across CS-US pairings. (**C**) Fear extinction decreased freezing across CS trial-block. In IL-NAc projection neurons, there were no differences following fear conditioning or fear extinction in (**H**) resting membrane potential, (**I**) action potential threshold, or (**J**) rheobase. (**G**) There was a significant reduction in the number of action potentials fired in the fear conditioning group when recordings were performed at -70mV, but was not present when performed at resting membrane potential. In PL-NAc projections neurons, there were no differences in (**H**) resting membrane potential, (**I**) action potential threshold, (**J**) rheobase, or (**K**) action potential number with increasing current injection magnitude. All data presented are mean ± SEM. N=4 mice per group, n=6-9 cells per group, per brain region. * denotes p<0.05.

Recording electrodes (3-5MΩ) were pulled from thin-walled borosilicate glass capillaries with a Flaming-Brown Micropipette Puller (Sutter Instruments, Novato, CA, USA). Recordings were performed only in cells expressing the fluorescent Retrobeads (pyramidal neurons of layer 2/3 and 5 of the PFC). Intrinsic neuronal excitability and current-injected firing was measured in current-clamp mode using electrodes filled with an intracellular recording solution containing (in mM) 135 K-Gluc, 5 NaCl, 2 MgCl_2_, 10 HEPES, 0.6 EGTA, 4 Na_2_ATP, and 0.4 Na_2_GTP. During recording, cells with access resistance greater than 30 MΩ, that had changes in access resistance greater than 10%, or that had action potentials that did not surpass zero were also excluded from the analysis.

Measurements of excitability were taken 5 minutes following the establishment of the whole-cell configuration. Intrinsic excitability was assessed via multiple measures: 1) resting membrane potential (RMP), 2) rheobase, defined as the minimum amount of current required to fire an action potential using a current ramp, 3) the action potential threshold, defined as the minimum voltage at which the neuron fired an action potential, and 4) the relationship between increasing steps of current and the action potentials fired using a voltage-current plot protocol (V-I plot). To control for differences in RMP, current-injection protocols were performed at both RMP and -70 mV. Signals were digitized at 10 kHz and filtered at 3 kHz using a Multiclamp 700B amplifier and analyzed using Clampfit 10.3 software (Molecular Devices, Sunnyvale, CA, USA).

### Chemogenetic pathway-specific inhibition

PFC outputs to the BLA were inhibited using DREADDs. Using the same stereotaxic coordinates and procedures described above, 0.3 μl of the retrograde Herpes Simplex Virus (HSV) containing Cre recombinase (HSV-hEF1α-mCherry-IRES-cre, MIT Vector Core, Cambridge, MA, USA) was bilaterally injected in the BLA. In addition, 0.3 μl of KOR-DREADD (AAV8-hSyn-DIO-HA-KORD-P2A-mCitrine, UNC Vector Core, Chapel Hill, NC, USA) or control vector (AAV5-eF1α-DIO-ChR2-eYFP, UNC Vector Core) was bilaterally injected into the PFC, focusing on the IL (for schematic, see **Figure 4A**). The injection coordinates were 1.85 mm anterior to bregma, 0.30 mm lateral to the midline and 2.65 mm ventral to the skull surface. Channel Rhodopsin 2 (ChR2) was used as a control as pilot experiments showed it resulted in a comparable degree of cell filling as the KOR-DREADD. It was selected on the basis that it was another non-endogenous transmembrane protein and would be insensitive to the administration of Sal-B.

**Figure 4:**
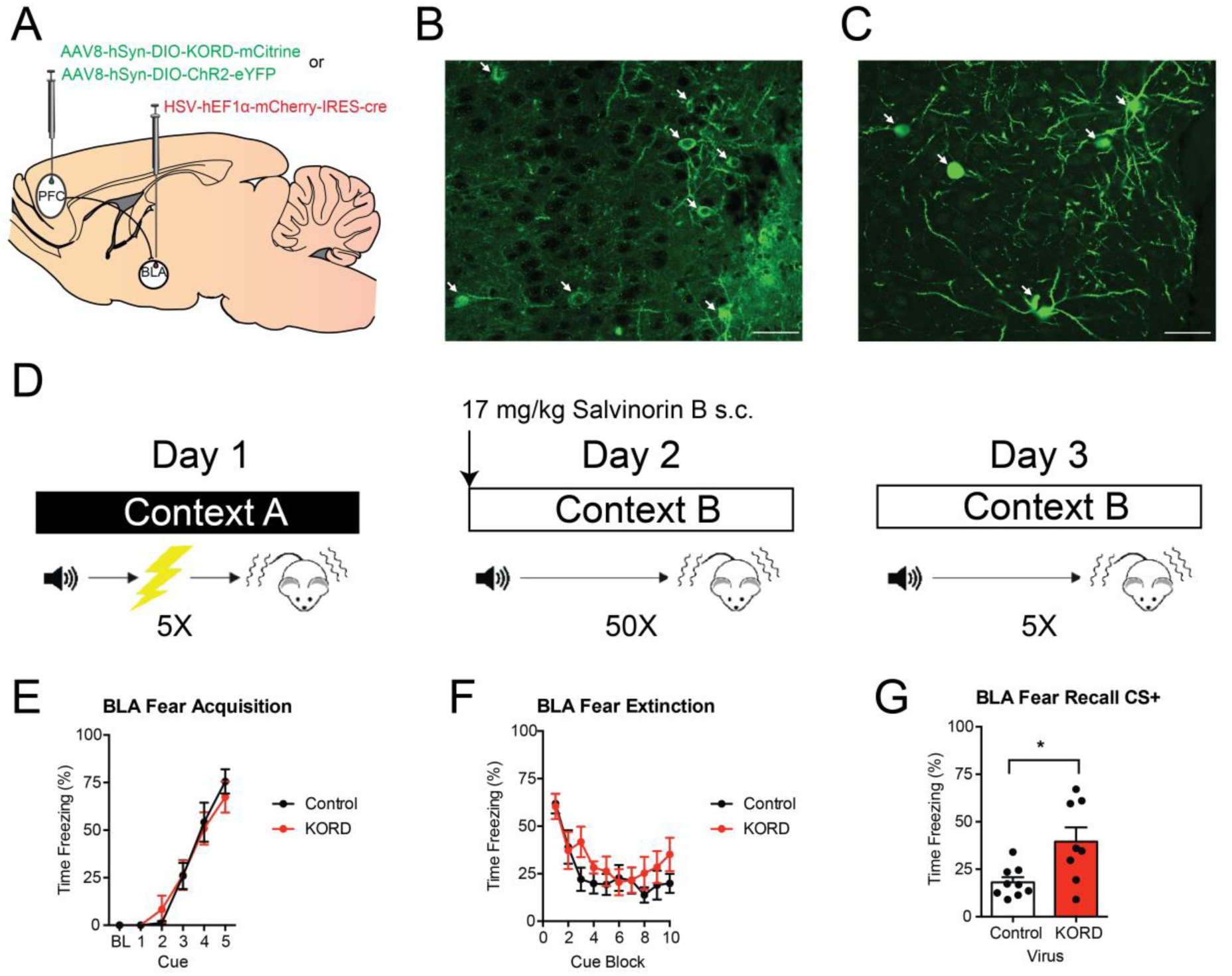
Chemogenetic inhibition of IL-BLA projections impairs long-term extinction memory formation. (**A**) HSV was injected into the BLA and KORD or a control virus was injected into the IL to selectively inhibit IL-BLA projections. Representative images of (**B**) KORD and (**C**) eYFP expression in the IL. (**D**) Mice were fear conditioned on day 1, then injected with Sal-B prior to extinction acquisition on day 2 and tested, drug-free, for extinction retrieval on day 3. Bottom row, PFC-BLA behavior cohort: (**E**) Fear conditioning increased freezing across CS-US pairings. (**F**) Fear extinction decreased freezing across CS presentations, irrespective of IL-BLA inhibition. (**G**) IL-BLA inhibition during extinction acquisition impaired extinction memory retrieval. All data presented are mean ± SEM. N=8-10 mice per group. * denotes p<0.05.

Six weeks following stereotaxic surgery to allow for virus expression, mice underwent fear conditioning, extinction acquisition and extinction retrieval, as described above (for schematic, see **Figure 4D**). Immediately before the fear extinction acquisition session, mice were subcutaneously injected with 17 mg/kg Salvinorin B (Sal-B) in DMSO at a 1 μl/g body weight injection volume using a 250 μl Hamilton Syringe (Hamilton Company, Reno, NV, USA). This dose was selected based on the initial characterization of the KOR DREADD (17). Mice were tested for extinction recall the following day, drug-free.

To verify virus expression at the completion of testing, mice were anesthetized, perfused and brains were processed, as described above. To visualize expression of HSV-mCherry-IRES-Cre in complex with the KOR-DREADD, a double immunohistochemistry labeling procedure directed against mCherry (to visualize Cre) and the HA epitope (to visualize KOR-DREADD) was performed using procedures previously described (17). Briefly, tissue sections were processed 3x in PBS, followed by 30 minutes of permeabilization in PBS with 0.3% Triton-X. Tissue sections were then blocked in 5% normal donkey serum and PBS in 0.3% Triton-X for 1 hour, and incubated with 1:500 rabbit anti-HA (cat# 3724S, RRID: AB_1549585, Cell Signaling, Danvers, MA, USA) and 1:500 mouse anti-mCherry (cat# ab65856, RRID: AB_1141717, Abcam, Cambridge, MA, USA) for 48 hours at 4 °C. For control brains, the same procedure was followed but substituting the HA primary antibody for an anti-GFP antibody recognizing eYFP (cat# GFP-1020, RRID: AB_10000240, Aves Labs, Tigard, OR, USA). Next, slices were washed 3x for 5 minutes in PBS with 0.3% Triton-X and incubated with 1:250 Alexa 488 donkey anti-rabbit (Cat# 711-545-152, RRID: AB_2313584, Jackson ImmunoResearch Labs, West Grove, PA) and Cy3 donkey anti-mouse secondary antibodies (Cat# 715-165-150, RRID: AB_2340813, Jackson ImmunoResearch Labs, West Grove, PA) for 24 hours at 4°C. Tissue sections were then washed 3x for 5 minutes in PBS with 0.3% Triton-X, followed by 2 x 5-minute washes with PBS. Slides were prepared with a Vectashield anti-fade mounting medium containing DAPI (Vector Labs, Burlingame, CA, USA).

Verification of virus injection site and expression was assessed using a wide-field epifluorescent microscope (BX-43, Olympus, Waltham, MA, USA) using a mouse stereotaxic atlas (Franklin & Paxinos, 2008). Mice with mis-targeted viral expression were excluded from the analysis. Representative images of virus expression were acquired a Zeiss 800 Laser Scanning Confocal Microscope (20x objective, NA 0.8) (Carl Zeiss, Jena, Germany). Images were cropped and annotated using Zeiss Zen 2 Blue Edition software (for example images, see **Figure 4B-C**).

### Statistical analysis

Groups were compared for RMP, rheobase and action potential threshold using a one-way analysis of variance (ANOVA), followed by Tukey’s *post hoc* tests. The relationship between current injection step and the number of action potentials fired was analyzed using 2-way repeated measures ANOVA, followed by Bonferroni *post hoc* tests. Groups were compared for freezing across CS-US pairing/CS-presentation during fear conditioning and extinction acquisition using 2-way repeated measures ANOVA followed by Tukey’s *post hoc* tests. Freezing during extinction retrieval was compared between groups using Student’s t-tests with Welch’s correction. Any outliers were detected and removed using Grubb’s outlier tests. All data are reported as mean ± standard error (SEM) and were analyzed with GraphPad Prism (GraphPad Software, San Diego, CA, USA).

## Results

### Separate IL and PL projections to BLA and NAc

We first sought to examine the relative weighting of IL and PL projections to the BLA and NAc, and quantify the degree to which these projections were distinct or overlapping. To accomplish this, we injected two different dye conjugates of the retrograde tracer, CTB, into the BLA and NAc and quantified the resulting overlap in the PFC (**Figure 1A-B**). Using this approach, we observed a significant proportion of BLA-labeled cells (38.2% of all cells counted) and NAc-labeled cells (58.4%), but only a small proportion of cells (3.4%) that were both BLA- and NAc-labeled (**Figure 1C-D**). These data suggest there are largely non-overlapped projections from the IL and PL to the two output regions. Importantly, we found that output neurons were located in both layer 2/3 and layer 5 in these regions, and thus our slice physiology experiments focused on output, rather than cell layer.

### Extinction increases the excitability of the IL-BLA, not PL-BLA, pathway

On the basis of our initial tracing data indicating separate populations of IL projections to BLA versus NAc, our next step was to use retrograde tracers to perform electrophysiological recordings from cells in a projection-specific manner. We injected fluorescent red Retrobeads (Lumafluor, Durham, NC, USA) into the BLA to test the differential effects of fear conditioning and extinction on the excitability of IL and PL output neurons (**Figure 2A**). Behaviorally, we confirmed that fear conditioning produced the expected increase in freezing across CS-US pairings that did not differ between the conditioned and to be extinguished groups (**Figure 2B**). Also as expected, repeated CS presentations in the extinction group led to decreased freezing to the cue during extinction acquisition (**Figure 2C**).

Slice physiological recordings revealed that IL-BLA cells were significantly more depolarized (more positive resting membrane potential) after fear conditioning, relative to test-naïve controls (F2, 21=4.23, p<0.01, *post hoc* test: p<0.05) (**Figure 2D**), and had a lower action potential threshold (potential at first-spike), relative to the extinction group (F2,21=4.78, p<0.05, *post hoc* test: p<0.05) (**Figure 2E**). Following extinction acquisition, IL-BLA cells had a significantly lower rheobase, as compared to naïve controls (F2,21=3.66, p<0.05, *post hoc* test: p<0.05) (**Figure 2F**). Additionally, extinction was associated with a significantly higher number of action potentials after current injection compared to both the naïve and conditioning only groups (F40,380=6.25, p<0.01) (**Figure 2G**). These various physiological changes were evident when neurons were held at RMP, but not at -70 mV, with the exception of the V-I plot differences, which were evident at both (data not shown). In contrast to the effects of extinction on IL-BLA projections, PL-BLA cells showed no group differences on any of the measures taken (**Figure 2H-K**), irrespective of whether recordings were performed at RMP or -70 mV. Together, these data indicate a preferential shift towards increased intrinsic excitability in IL-BLA cells following fear extinction, and suggest that this pathway is more likely to be activated after mice have underwent extinction.

### Extinction reverse fear-related plasticity in the IL-NAc pathway but does not alter PL-NAc

Our next step was to test whether the extinction-related changes in IL projections to the BLA were present or absent in IL and PL neurons projecting to the NAc. In these mice, fear conditioning produced an increase in freezing across CS presentations that did not differ between groups (**Figure 3B**). Moreover, extinction training resulted in a decrease in freezing to the CS during the extinction conditioning (**Figure 3C**). However, neither conditioning nor extinction produced any significant change in resting membrane potential, action potential threshold, or rheobase whether recorded at RMP (**Figure 3D-F**) or at -70 mV (data not shown). At -70 mV (but not RMP, data not shown), action potential firing increased with current injection magnitude in a manner that was lower in the fear conditioned group, relative to naïve controls, but not the extinction group (current x group interaction: F40,19=1.68, p<0.05) (**Figure 3G**). When we recorded from PL-NAc cells, no significant group differences were found in any measure of intrinsic excitability (**Figure 3H-K**). Together, these recording results show that the fear conditioning leads to a suppression of firing in IL neurons that project to the NAc, which is reversed following extinction.

### Inhibiting PFC-BLA projections impairs extinction memory recall

Thus far, our anatomical and physiological data implicate the IL-BLA pathway in extinction, but do not provide the kind of causal evidence for such a role that has been demonstrated by other studies. To test this hypothesis, we used a dual virus chemogenetic approach to inhibit PFC outputs to the BLA (**Figure 4A**). Infusion of HSV-mCherry-IRES-Cre into the output region resulted in dense labeling in the PFC in KOR-DREADD (**Figure 4B**) and ChR2-eYFP animals (**Figure 4C**).

In the PFC-BLA experiment, fear conditioning significantly increased freezing across CS-US pairings (F4,64=60.68, p<0.01), irrespective of virus group (**Figure 4E**). Following Sal-B injection prior to extinction acquisition, freezing significantly decreased across trial-blocks (F9,144= 13.48, p<0.01) in a manner that did not differ between virus groups (**Figure 4F**). However, on drug-free extinction retrieval the following day, freezing was significantly higher in the KORD group than in controls (t(8.77)=2.72 p=0.02) (**Figure 4G**).

## Discussion

The main findings of the current study were that IL neurons projecting to the BLA exhibited significant changes in intrinsic excitability following fear extinction acquisition, and that selectively chemogenetically inhibiting this pathway was sufficient to disrupt extinction memory formation. The current results substantiate much of the findings of earlier studies that establish a critical contribution of the IL to fear extinction.

Employing techniques including immediate-early gene and molecular mapping, *in vivo* single-unit recordings, lesions, and IL-targeted pharmacological and optogenetic manipulations, a considerable body of work has shown that IL activity positively predicts successful fear extinction, whereas functionally disrupting this region impairs extinction (1, 4-6). Of particular relevance to the current findings, Porter and colleagues have used *ex vivo* electrophysiological recordings to demonstrate changes in plasticity and the intrinsic excitability of IL neurons as a result of fear extinction (18-20). Taken together with the current finding that there are similar changes in excitability in IL-BLA cells, these electrophysiological data suggest that extinction recruits IL neurons projecting to the BLA. Indeed, this conclusion fits well with recent studies using neuronal tracing, electrophysiology and optogenetics to show that IL inputs directly innervate BLA neurons (9, 21) to produce downstream changes in BLA plasticity (8) that instruct extinction memory formation (7, 22).

Another notable contribution of the current study lies in clarifying which outputs are not recruited by fear extinction. By employing a dual retrograde tracer strategy, we observed that projections to the BLA are largely distinct from NAc projections, and may explain why fear extinction results in different electrophysiological responses in these populations. In this context, it has recently been shown that different outputs of the PFC become activated by positive and negative emotional experiences (23). Given the well-established role for PFC-NAc projections in the extinction of cued drug seeking (24, 25), it is reasonable to posit that this output is more actively involved in the extinction of appetitive behaviors. It does bear noting that the retrograde tracing technique employed here did not label the entire BLA and NAc and thus, the number of CTB positive cells in the PFC is likely an underestimation of the number of outputs to either region. However, this experiment is not intended to provide an absolute measure of the number of projection neurons, but rather a measure of the relative size and overlap of these two populations. The high degree of CTB labeling in layer 2/3 cortex is surprising, as layer 5 is canonically thought as the output layer of cortex. Previous monosynaptic rabies tracing studies have, however, noted there are direct connections from layer 2/3 cortex to the striatum (26), and this preferential labeling of layer 2/3 cortex has been observed in previous retrograde tracing studies of the PFC-BLA pathway (27).

There are several studies that, at first glance may seem to be odds with our findings. Specifically, when performing non-projection specific recordings in the IL, Santini and colleagues report a decrease in IL excitability following conditioning which is reversed by extinction (only to naïve levels) (17). Additionally, Cruz and colleagues have reported changes in IL intrinsic excitability exclusively starting at extinction consolidation (specifically, 4 hours after extinction), which are absent immediately after extinction (18). However, these studies were performed in Sprague-Dawley rats, and thus it is possible that these differences are attributable to the difference in species. Alternatively, as the current study used retrograde tracers to prospectively identify outputs to the BLA and NAc for recordings, the differences observed may be due to the specific population of neurons sampled. Consistent with the findings of these two prior studies, we found decreased excitability in the IL-NAc pathway following fear conditioning that returns to naïve levels following extinction. Given the fact that we observed slightly greater numbers of IL cells projecting to the NAc than the BLA, it is possible that the cells sampled by Santini et al. may have been weighted more heavily towards this population. This suppression and restoration of excitability in IL-NAc neurons may be consistent with a model in which fear extinction restores the excitability of an appetitive pathway. In addition to changes in excitability in the PFC, previous studies have noted that fear extinction results in decreased synaptic efficacy of PFC inputs to the BLA (21, 27). As we find increased excitability of PFC-BLA neurons following extinction, these findings suggest that it is possible that there is compartmentalization of plasticity events in cell body and terminals. This will be an important question for future studies.

The current chemogenetic data provide an important confirmation and extension of recent work showing that optogenetically exciting IL projection neurons in the BLA or BMA promotes extinction, whereas silencing these cells impairs extinction (7, 10). Through using a dual-virus system in which an HSV virus was injected directly into the BLA, taken up by terminals therein and retrogradely transported to the IL for recombination with the KOR-DREADD virus, the current approach mitigates potential effects on BLA fibers of passage and demonstrates that IL neurons terminating in BLA are necessary for extinction memory formation. However, there are several important technical caveats to our findings. First, because we gave Sal-B systemically, it is possible that the PFC neurons that project to the amygdala also project to other regions, and it is via actions in these other regions that disrupt extinction. Unfortunately, the solubility of the KORD agonist precludes local infusion. In addition, we did not include a vehicle control group. However, we do not consider this a major concern, as the concentration of Sal-B we used does not cause changes in locomotion or have any anti-nociceptive properties (17, 28). Another methodological factor to point out is that there was an age difference in the mice used for the DREADD experiments (14 weeks) versus the mice used in the electrophysiology experiments (11 weeks). Given the behavior of the mice in the two experiments, these age differences are not likely to have impacted our conclusions. Finally, the use of the agonist Sal-B, which has a relatively short active duration (45-90 minutes) (17), allowed the DREADD activation to be timed so that chemogenetic inhibition roughly corresponded with the length of the extinction session. However, it is possible that the results observed here are from continued inhibition of IL during extinction memory consolidation. Notably though, these data replicate the findings of Quirk and collegues (29) whereby brief optogenetic inhibition of IL during extinction learning, but not extinction retrieval, impaired the expression of the extinction memory. Other groups have found opposing results though(30), highlighting the need for future studies to more precisely refine the time course during IL is active during extinction learning and retrieval.

A number of interesting questions remain to be addressed in future work. One important issue to clarify is the precise downstream targets of the IL inputs that subserve extinction. This includes ascertaining the BLA neuronal populations and interneuron cell types that are directly innervated by the IL, as well as certain cell populations that neighbor, are tightly coupled to the BLA and have been implicated in extinction, particularly the intercalated cell clusters (8, 31-35). In addition, recent data from Goosens and colleagues suggests that extinction can specifically recruit BLA-NAc circuits to suppress fear reinstatement (13). Moreover, optogenetic stimulation of BLA terminals in the NAc resulted in increased c-fos labeling in the IL, raising the possibility of an IL-BLA-NAc feedback circuit that promotes extinction. There is then the question of the origin of the inputs to the IL that drive extinction-related increases in neuronal excitability. A good candidate for this is the BLA itself, based on recent evidence that BLA-IL and BLA-PL inputs exhibit functional changes that correlate well with extinction and fear, respectively (11, 12, 36). Thus, in light of the current findings and related data addressing the IL to BLA pathway, there would appear to be a bidirectional corticoamygdala circuit underlying extinction, rather than a simple ‘top-down’ (cortical to limbic) route that had been initially proposed.

In summary, the results of the current study offer further support for the importance of a neuronal circuit from the IL to the BLA in Pavlovian fear extinction. By employing a variety of approaches, we were able to show that IL, but not PL, projection neurons innervating the BLA displayed an increase in the intrinsic excitability as a result of fear extinction. In contrast, an anatomically distinct population of IL neurons projecting to the NAc did not exhibit such changes, however there appeared to be an extinction driven reversal of plasticity, in keeping with results from Porter and colleagues. At the behavioral level, using DREADDs to selectively inhibit PFC-BLA neurons, we demonstrated the necessity of this pathway for extinction memory formation. Collectively, these findings build on a growing literature establishing the critical contribution of the IL-BLA neural circuit to fear extinction. Given evidence of extinction-related aberrations in an analogous circuit in patients with trauma-related disorders (1, 37), these current results could help to illuminate understanding of the pathophysiology and routes to treating these conditions.

## Acknowledgements

We would like to thank Hee-Jung Jo and Maria Luisa Torruella-Suarez for assistance with histology and immunohistochemistry.

